## Supplementary material for "TaxiBGC: a Taxonomy-guided Approach for Profiling Experimentally Characterized Microbial Biosynthetic Gene Clusters and Secondary Metabolite Production Potential in Metagenomes": Figure S1

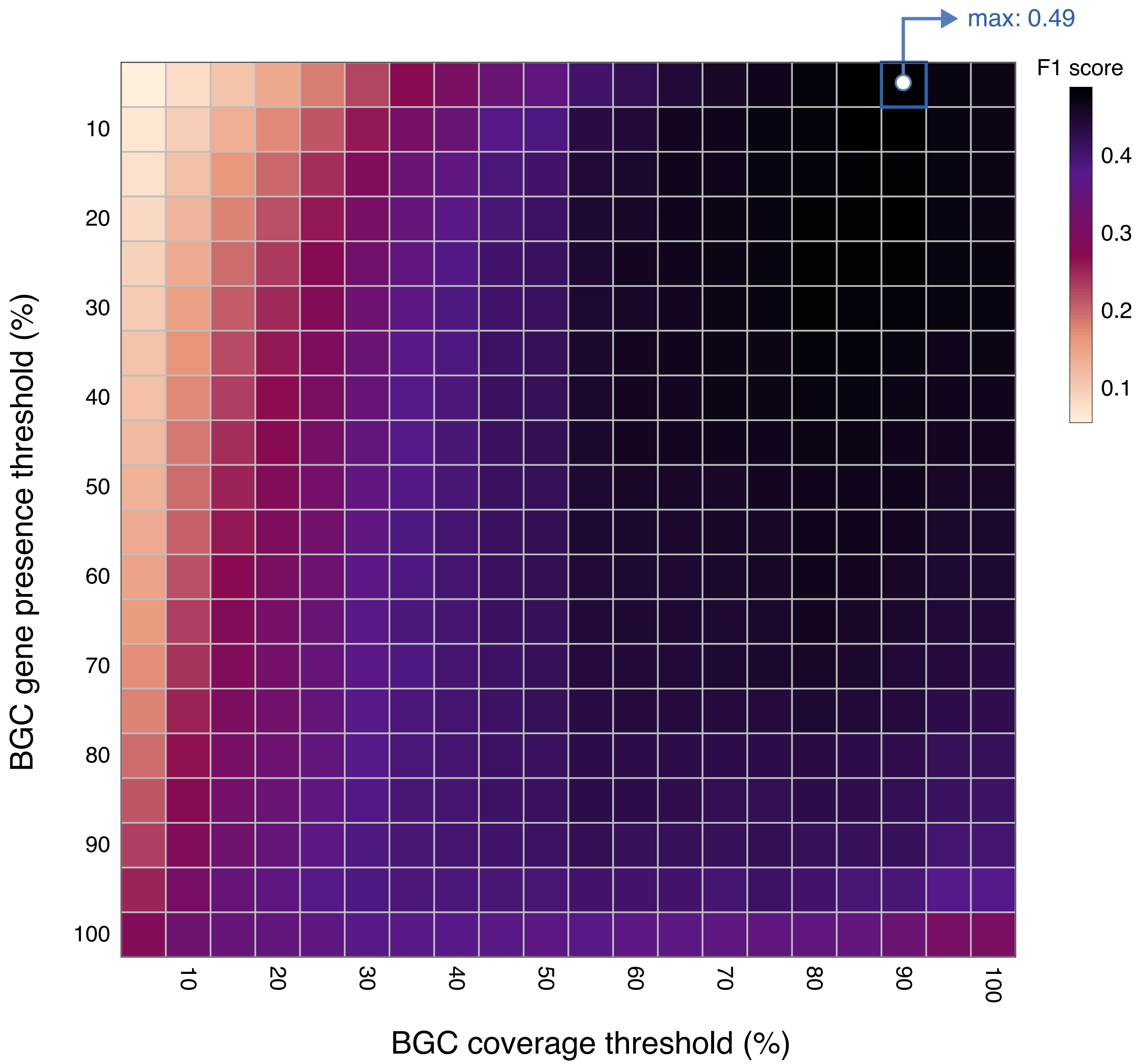


**Figure S1. Predictive performance of the direct BGC detection approach based on two different thresholds.** The direct BGC detection method was used to predict BGCs in all 125 simulated metagenomes of mock microbial communities. We used 400 total pairwise combinations of BGC gene presence (5–100% with an interval of 5%) and BGC coverage (5–100% with an interval of 5%) thresholds. The best overall accuracy for predicting BGCs in simulated metagenomes was achieved with a minimum BGC gene presence and BGC coverage of 5% and 90%, respectively (mean F1 score: 0.49).
