## Supplementary material for "TaxiBGC: a Taxonomy-guided Approach for Profiling Experimentally Characterized Microbial Biosynthetic Gene Clusters and Secondary Metabolite Production Potential in Metagenomes": Figure S2

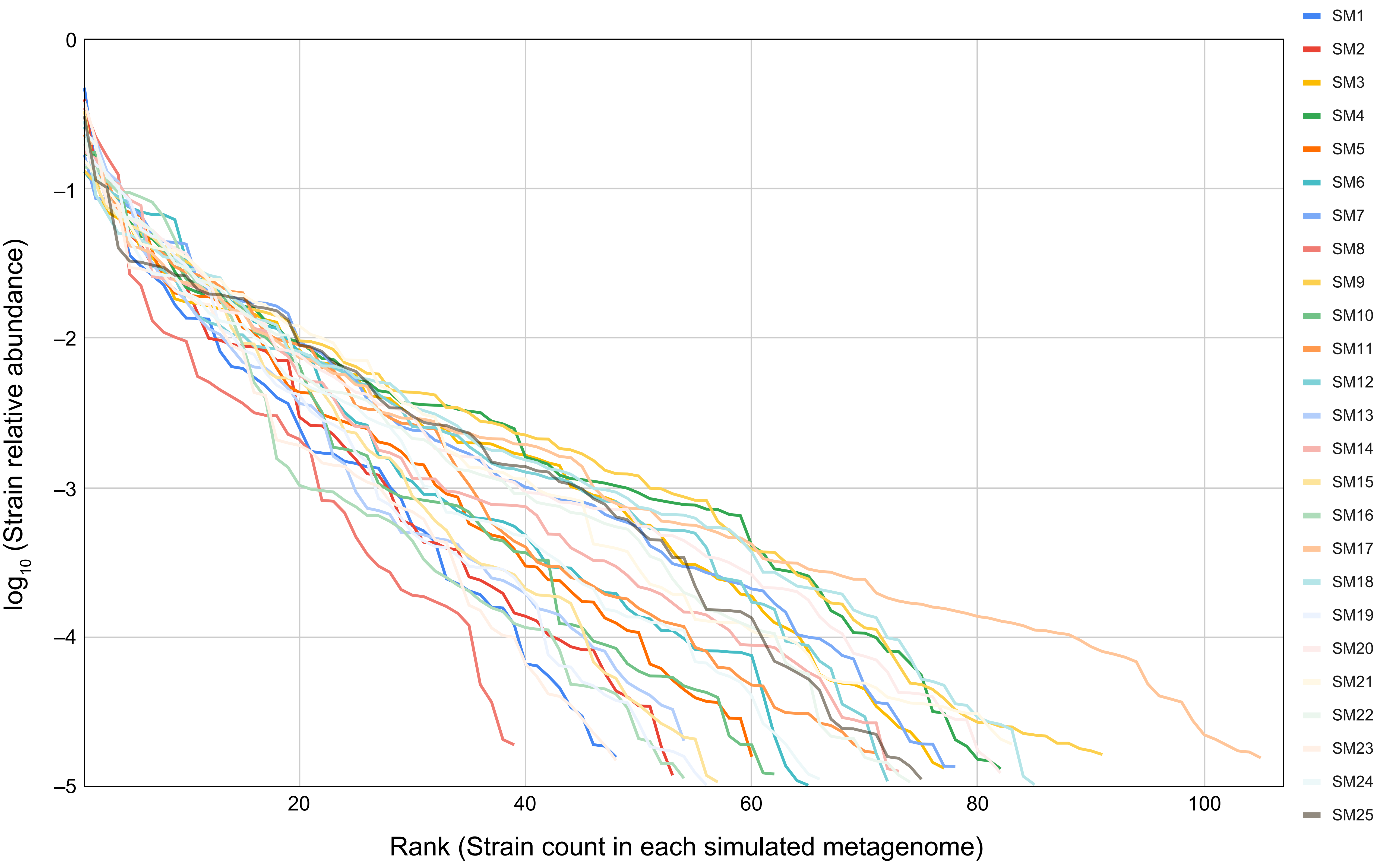


**Figure S2. Relative abundances of strains in each simulated metagenome.** The rank-plot shows (in descending order of magnitude) the relative abundances of all strains in each simulated metagenome. The mock communities used to construct simulated metagenomes are composed of 39–105 unique strains, resulting in 39–97 unique species. SM, simulated metagenome.
